# *FAST^m^C:* a suite of predictive models for non-reference-based estimations of DNA methylation

**DOI:** 10.1101/029496

**Authors:** Adam J. Bewick, Brigitte T. Hofmesiter, Kevin Lee, Xiaoyu Zhang, Dave W. Hall, Robert J. Schmitz

**Author notes:** **CORRESPONDENCE** Robert J. Schmitz.

## Abstract

We describe a suite of predictive models, coined *FAST^m^C,* for non-reference, cost-effective exploration and comparative analysis of context-specific DNA methylation levels. Accurate estimations of true DNA methylation levels can be obtained from as few as several thousand short-reads generated from whole genome bisulfite sequencing. These models make high-resolution time course or developmental, and large diversity studies practical regardless of species, genome size and availability of a reference genome.

## BACKGROUND

Advances in high-throughput sequencing has allowed for single-base resolution analysis of DNA methylation at cytosines across an entire genome. This was first applied to the model plant *Arabidopsis thaliana* [1],[2] and, since then, has been applied to numerous species, including protists, fungi, insects, anthozoa, tunicates, fish, and mammals [3]-[5]. Currently, DNA methylation is profiled genome-wide by deep, whole-genome bisulfite sequencing (WGBS). The use of a reference genome is essential to inform the methylation status at each cytosine reference position, where a thymine *in lieu* of cytosine indicates an unmethylated cystosine [6]. Thus, absence of a reference genome has prevented rapid, genome-wide analysis of DNA methylation for the majority of known species, and is cost-prohibitive for high-resolution developmental or time-course studies in species with large genomes. To date, several methods exist to accommodate the challenges assocaited with non-reference based analysis of DNA methylation, but lack cytosine context sequence specificty [7]-[9].

Here we present *FAST^m^C,* a suite of predictive models that can be used to estimate genome-wide DNA methylation levels at all cytosine sequence contexts without the use of a reference genome. These models assumed a relationship between DNA methylation levels calculated from alignment of WGBS reads to a reference genome (target; *m*) and from direct assessment from raw WGBS reads (i.e., no alignment to a reference genome) (estimator; 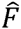). Methylation levels are calculated as the proportion of methylated cytosines to the total number of possible methylated cytosines. The difference between the two variables exists at unmethylated cytosines; the estimator value includes unmethylated cytosines and true thymines when calculating the DNA methylation level. Estimator DNA methylation levels were compared to target levels to determine a relationship, and the strength of which, to confidently predict/extrapolate genome-wide DNA methylation levels for any sample regardless of the availability of a reference genome.

Using publicly available data, for species with reference genomes, target and estimator DNA methylation levels for 44 species were used to construct models capable of predicting genome-wide levels of DNA methylation for species without a sequenced genome. Using additional publicly available data from mutants and cell-types known to be different from wild-type samples, we discuss the sensitivity, robustness and utility of the models in terms of CpG DNA methylation, followed by plant- (CHG and CHH) and mammal-specifc (CH) DNA methylation.

## RESULTS AND DISCUSSION

*FAST^m^C* is able to detect intraspecific differences in DNA methylation (Fig. 1). In the plant *A. thaliana,* mutants exist that are defective for enzymes that are required for maintenance of CpG DNA methylation – *met1, met1+cmt3,* and *vim1+vim2+vim3* – as they have reduced CpG methylation levels compared to wild type [10]. Also, several mutant genotypes for *met1* show different degrees of loss of CpG DNA methylation compared to each other: (i) An original *met1* mutant genotype (high loss); (ii) A *met1* heterozygous mutant genotype *(met1 +/*−) (intermediate loss); and (iii) A recovered genotype *(MET1* +/+) from a *MET1* +/+ and *met1* +/− backcross. The recovered *MET1* +/+ is wild-type for MET1 function but has lost CpG methylation in some regions of the genome (low loss). *FAST^m^C* is able to capture the differences between these maintenance methyltransferases (Fig. 1A). Additionally, the slight (~3%) difference between *MET1 +/+* and the *met1* +/− mutant can be distinguished, demonstraing the sensitivity of *FAST^m^C* (Fig. 1A).

**Figure 1.**
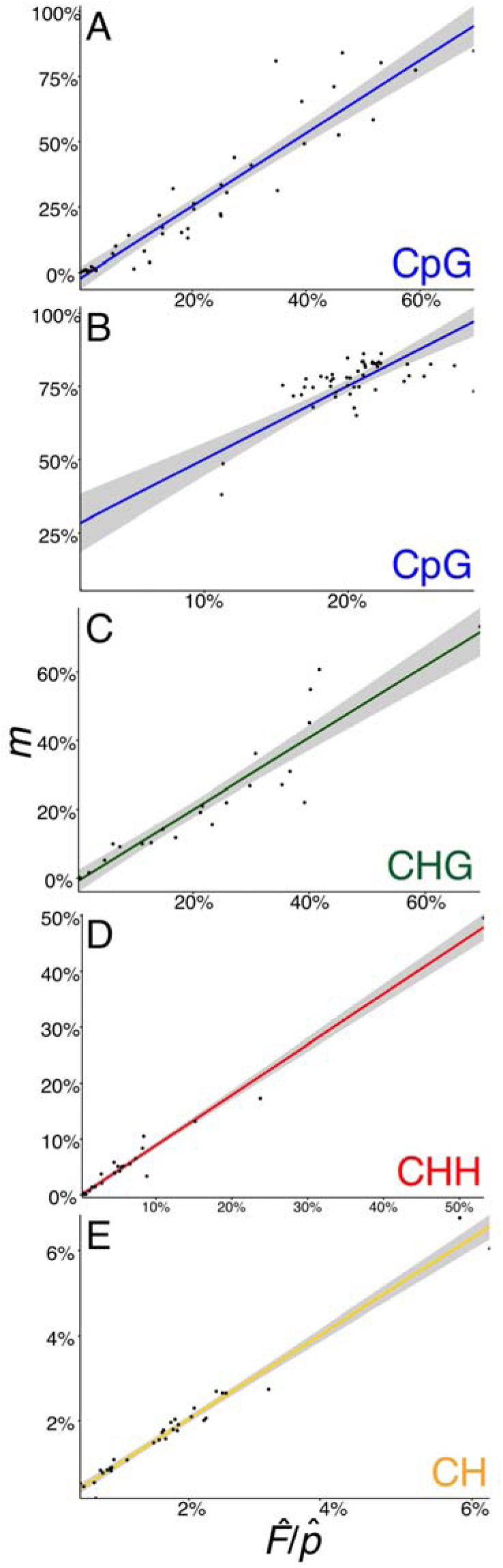
Detection of intraspecific DNA methylation levels by *FAST^m^C.* Generalized linear models (GLMs) for estimator 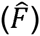 versus target (*m*) CpG, CHG, CHH, and CH DNA methylation levels using 10,000 reads corrected for estimated GC content 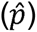 (A-E). Differences between *A. thaliana* mutants can be detected (A, C, and D). Also, differences of CpG DNA methylation between mutants, cell-types and tissues in *M. musculus* can be differentiated by *FAST^m^C* (B). Finally, increasing CH methylation through brain development is captured with *FAST^m^C* (E). Shaded area represents the 95% confidence interval.

In mammals, epigenetic reprogramming, including CpG demethylation, is required to erase DNA methylation imprints and epimutations established in the previous generation [11]. Following demethylation, DNA methylation patterns are re-established at imprinted loci and transposable elements (TEs) during gametogenesis by the *de novo* methyltransferases DNMT3A and a non-catalytic paralogue, DNMT3-like (DNMT3L) (reviewed by [12]). The reductions in CpG DNA methylation caused by epigenetic reprogamming in primordial germ cells (PGCs) or by mutations in DNMT3L (*dnmt3L*) compared to somatic tissues are captured by *FAST^m^C* (Fig. 1B) [13]-[15]. Additionally, increased levels of CpG DNA methylation in the brain (e.g., *NeuN+ and glia* cells) [16] can be differentiated from other somatic tissues (Fig. 1B; Suppl. Table 1) [17]. Overall, as demonstrated in *A. thaliana* and *M. musculus, FAST^m^C* can be used to accurately detect intrapecific differences of DNA methylation levels at CpG sites (Fig. 1A and B).

We determined natural interspecific variation of DNA methylation at CpG sites across 44 different species (Fig. 2A). However, unlike intraspecific comparisons between mutants or cell-types, nucleotide biases, such as genomic GC content differences, can over- or underestimate the estimator value for the CpG sequence contexts. The estimator (equation 2 of Methods) is estimating the product of the methylation frequency of CpG sites and the GC content of the genome, and are thus confounded. This bias can be overcome in all species investigated but mammals (*H. sapiens, M. musculus,* and *C. l. familiaris*) by dividing the estimator value by an average GC content of the genome, which corrects the relationship between target and estimator to ~1:1. GC content can be approximately estimated from WGBS reads (see Methods) or additional genomic sequence data − 10,000, 50 base pairs (bp) reads (500,000bp) – can be used to directly estimate GC content (Suppl. Table 1).

**Figure 2.**
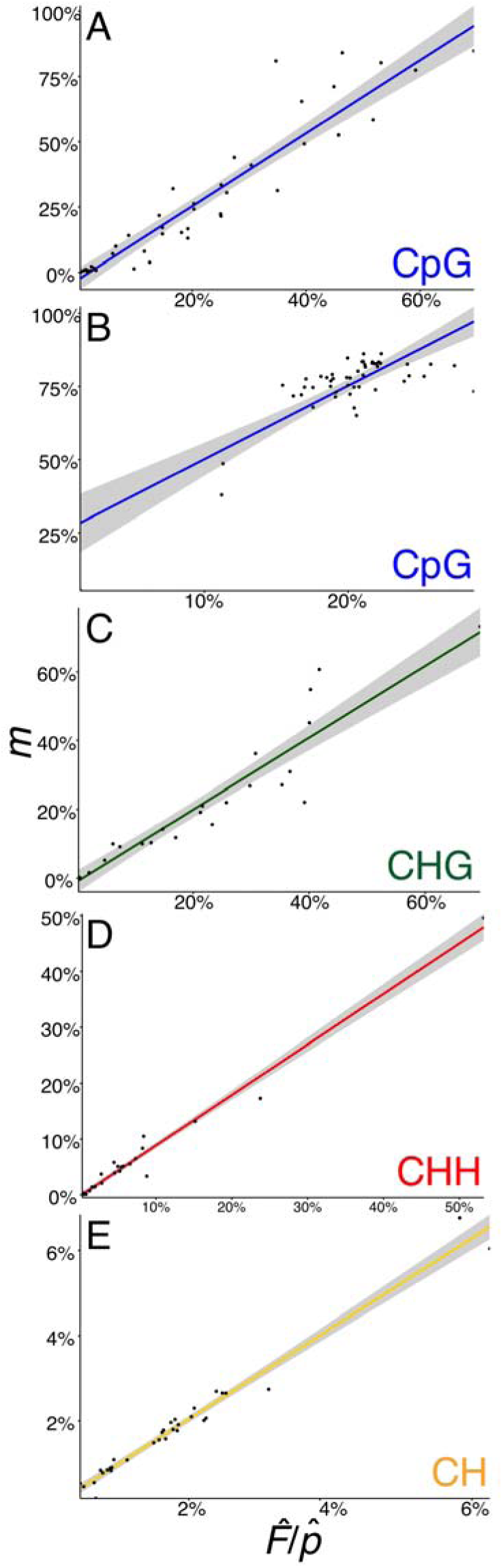
Detection of interspecific DNA methylation levels by *FAST^m^C.* Generalized linear models (GLMs) for estimator 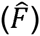 versus target (*m*) CpG (A-B), CHG (C), CHH (D), and CH (E) DNA methylation levels using 10,000 reads corrected for estimated GC content 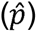. Species included in each plot can be found in Suppl. Table 1. Shaded area represents the 95% confidence interval.

Nucleotide biases in genomes – such as the depletion of CpG dinucleotides to localized “CpG islands” in mammalian genomes – may interfere when estimating 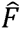. CpG dinucleotides can be directly measured from 10,000, 50 bp genomic sequencing reads (Suppl. Table 1), and this can then be used to directly calculate the proportion of target sites that are methylated, *m,* using the frequency of intact target sites, e.g., CpG, that remain in the bisulfite sequencing data. These are sites that were methylated and thus escaped C to T conversion. Accommodating for nucleotide biases in mammalian genomes does not improve assessment of DNA methylation levels by *FAST^m^C* (Suppl. Table 1). However, treating mammals separately from other species with CpG DNA methylation (i.e., phylogenetic correction) produces an improved, mammal-specific model with similar accuracy – measured as the Mean Absolute Percentage Error (MAPE) – to the remaining species (Suppl. Table 1). Additionally, only a modest increase in model improvement was observed for non-mammalian species (Suppl. Table 1). Overall, GC content correction 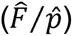 and treating mammalians species separately improves model accuracy without introducing additional genomic sequencing data.

*FAST^m^C* also tolerates high contamination and error rates associated with sodium bisulfite conversion. We used *A. thaliana met1* mutants generated by [10], which show minor (~3%) to large (~14%) differences in CpG DNA methyaltion compared to the wild-type *A. thaliana.* By artificially introducing un-methylated chloroplast reads to 10,000 reads to *met1* and *met1 +/*− mutant genotypes, and *MET1* +/+ and *A. thaliana* wild-type genotypes, we were able to demonstrate that a ~3% difference in DNA methylation can still be detected with <10% chloroplast contamination, and a difference of 13-14% with 40-50% chloroplast contamination (Suppl. Table 1). Similarly, nonconversion rates >3% still allow for detection of differences between samples (Suppl. Table 1). It should be noted that the *met1* mutants and *A. thaliana* samples had nonconversion rates of 0.50%, 0.82%, 1.86%, and 0.56% for *met1, met1 +/*−, *MET1 +/+,* and wild-type *A. thaliana,* respectively. The artifically introduced error rates are extremely high, but possible. For example, <1% of reads typically map to the chloroplast genome, and nonconversion rates are typically <2% (data not shown). However, it is recommended that Lambda DNA be sequenced for each batch of WGBS libraries prepared to estimate the rate of sodium bisulfite non-conversion. Reducing technical error is especially important for identifying differences between species with small amounts of or no DNA methylation like insects (Suppl. Table 1). Regardless, the *FAST^m^C* method is robust as it is able to tolerate technical and biological contamination.

The number of short reads (≥30 bp) required to make accurate estimations is low, and we have determined that a few thousand reads produce high-confidence estimates of genome-wide methylation levels (Suppl. Fig. 1). Hence, these models can be used to accurately, and cost-effectively, identify differences of DNA methylation levels for any species regardless of the availability of a reference genome assembly.

Non-CpG DNA methylation can also be confidently predicted within and between species using *FAST^m^C.* In *A. thaliana,* the majority of DNA methylation at CHG sites is maintained by chromomethylase CMT3 through a reinforcing loop with H3K9me2 methylation catalyzed by the KRYPTONITE (KYP)/SUVH4 protein [18]-[20]. Similarly to MET1, mutations in CMT3 causes reductions in CHG DNA methylation [10], which are accurately detected by *FAST^m^C* (Fig. 1C). Also, in *A. thaliana,* cell-type specific levels of CHH DNA methylation in the sperm cell (SC) (i.e., hypo-CHH DNA methylation) and vegetative nucleus (VN) (i.e., hyper-CHH DNA methylation), and depletion of CHH DNA methylation in mutants in the *de novo* DNA methylation pathway (e.g., the DNA-dependent RNA polymerase, NRPD1) were recapitulated (Fig. 1D) [21],[10].

In mammals, non-CpG DNA methylation can be found at CH sites. Work by [16] has demonstrated the overall increase of CH DNA methylation during brain development in *M. musculus* and *Homo spaiens. FAST^m^C* was able to capture the overall trend of increasing CH methylation through brain development in *H. sapiens* (Fig. 1E). Furthermore, despite only small differences in brain CH methylation in the intervals from 2 years to 5 years (0.068%), and from 55 years to 64 years (0.062%) of age, the *FAST^m^C* model accurately detected these changes (Fig. 1E) [16].

## CONCLUSIONS

We propose several models, which capture the variation of, and can accurately predict, genome-wide DNA methylation levels between species to represent *FAST^m^C* and can be found at http://fastmc.genetics.uga.edu. Additionaly, the web-based interface makes *FAST^m^C* universally accessible, and models will be continuouslly updated when new whole genome and methylome data is analyzed and becomes available. Although genome content biases interfere with the accuracy of *FAST^m^C,* treating mammalian species separately for CpG DNA methylation overcame this obstacle. *FAST^m^C* makes practical previously intractable studies (e.g. high-resolution time course, developmental, and large diversity panels) regardless of species, genome size and availability of a reference genome. Furthermore, these models will greatly contribute to high-resolution screening of either developmental- or environmental-induced epigenomic reprogramming events. *FAST^m^C* is a suite of powerful models that can aid researchers to make better investments in more comprehensive, fruitful studies.

## METHODS

Whole genome bisulfite sequencing (WGBS) data was downloaded from the Short Read Archive (SRA)/Gene Expression Omnibus (GEO) or sequenced in-house (Suppl. Table 1). WGBS data was aligned using methods described in [22] to generate “allC” files. The allC files were used to determine target DNA methylation levels, and can be downloaded from GEO under accession number GSE72155. Prior to estimation of predictor DNA methylation levels, WGBS data was trimmed of adaptor sequences using Cutadapt v1.9 [23], end-trimmed using Trimmomatic [24], and quality filtered using FASTX-toolkit (http://hannonlab.cshl.edu/fastx_toolkit/). Reads of at least 30 base pairs (bp) in length with ≥20% of nucleotides having a quality score ≥75% were retained. Random sampling without replacement was performed with increasing fold-change from 1-10^5^ reads using the program fastq-tools (http://homes.cs.washington.edu/~dcjones/fastq-tools/). Custom Perl scripts were used to sum the number of C^m^ and C^?^ sites for each randomly sampled read, and subsequently to estimate the predictor DNA methylation level at CpG, CHG, CHH, and CH sites (Suppl. Table 1).

Predictive modeling is used to find the mathematical relation between a target, (dependent variable) and various estimators (independent variables); subsequent values of an estimator(s) are used to predict the target variable using the established mathematical relationship between them. The goal of the *FAST^m^C* models were to predict reference-based (target) from non-reference-based (estimator) DNA methylation levels. These models assume that in MethylC-Seq data [6]: (i) all cytosines at CpG, CHG, CHH, and CH sites are methylated. (ii) all thymines at TpG, THG, THH, and TH sites are converted unmethylated cytosines or true thymines, and (iii) all nucleotides are randomly distributed in the genome. Our goal is to estimate the proportion of Cs in potential target sites that are in fact methylated, *m,* which is

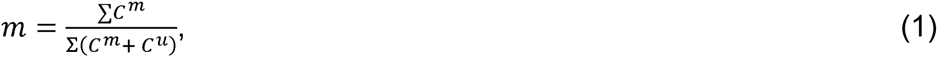

where ∑*C^m^* and ∑*C^u^* are the total number of methylated and unmethylated target sites in the genome, respectively. Since *m* is unknown, we use an estimator, 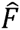, which is obtained from the bisulfite sequencing data:

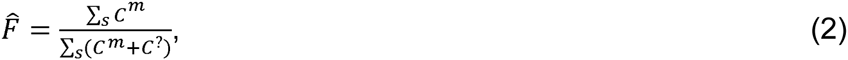

where ∑*_s_ C^m^* is the total number of methylated target sites in the sample and ∑*_s_ C*^?^ is the sum of unmethylated target sites plus sites that are equivelent to unmethylated target sites after bisulfite sequencing in the sample, e.g. all TG dinucleotides in the case of CpG methylation. With our assumptions, it is straightforward to show that for CpG methylation, the expected value of 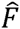 is *mp.* Thus, 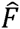 divided by the estimated genomic GC content, 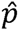, is an estimate of *m.* We estimate GC content from the frequencies of G nucleotides in the sample because these sites are unaffected from bisulfite treatment. Estimates of GC content from WGBS reads are on average within 4.56% ± 3.52% standard deviations of the true GC content. For the other three targets of methylation (CH, CHH and CHG), it can be easily shown that 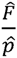 is also equal to *m. FAST^m^C* calculates 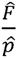 from a whole genome bisulfite sample and uses it to estimate *m,* the fraction of Cs that are methylated.

Violation of the assumptions can cause inaccuracies in estimating 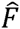. We discuss some of these vioations in the results section. In addition, we note that when additional genomic short read data (≥500,000 bp) is available, the frequency of the target site in the genome, e.g., the GC content and frequency of CpG dinucleotides, can be directly measured. This can then be used to directly calculate the proportion of target sites that are methylated, *m,* using the frequency of intact target sites, e.g., CpG, that remain in the bisulfite genome data. These are sites that were methylated and thus escaped C to T conversion.

## AVAILABILITY OF SUPPORTING DATA

All data used in this study can be found on the Short Read Archive (SRA)/Gene Expression Omnibus (GEO) webpages. Accession identifiers can be found in Suppl. Table 1.

## LIST OF ABBREVIATIONS

Whole-Genome Bisulfite Sequencing (WGBS), Primordial Germ Cells (PGCs), Base Pairs (bp), Short Read Archive (SRA), Gene Expression Omnibus (GEO), Mean Absolute Percentage Error (MAPE), Sperm Cell (SC), Vegetative Nucleus (VN)

## AUTHORS’ CONTRIBUTIONS

All authors contributed equally to this work.

## ACKNOWLEDGEMENTS

We would like to thank Nathan Springer for critical comments on this manuscript. Also, we would like to thank David Brown for webpage setup. The study was funded by grants from the National Science Foundation (MCB-1339194) and the National Institutes of Health (R00GM100000) to RJS.

## REFERENCES

1. Cokus, S. J., Feng, S., Zhang, X., Chen, Z., Merriman, B., Haudenschild, C. D., Pradhan, S., Nelson, S. F., Pellegrini, M., and Jacobsen, S. E. Shotgun bisulfite sequencing of the *Arabidopsis* genome reveals DNA methylation patterning. Nature 2008;452:215–219.

2. Lister, R., O’Malley, R. C., Tonti-Filippini, J., Gregory, B. D., Berry, C. C., Millar, A. H., and Ecker, J. R. Highly integrated single-base resolution maps of the epigenome in *Arabidopsis*. Cell 2008;133:523–536.

3. Lister, R., Pelizzola, M., Dowen, R. H., Hawkins, R. D., Hon, G., Tonti-Filippini, J., Nery, J. R., Lee, L., Ye, Z., Ngo, Q.-M., Edsall, L., Antosiewicz-Bourget, J., Stewart, R., Ruotti, V., Millar, A. H., Thomson, J. A., Ren, B., and Ecker, J. R. Human DNA methylomes at base resolution show widespread epigenomic differences. Nature 2009;462:315–322.

4. Feng, S., Cokus, S. J., Zhang, X., Chen, P.-Y., Bostick, M., Goll, M. G., Hetzel, J., Jain, J., Strauss, S. H., Halpern, M. E., Ukomadu, C., Sadler, K. C., Pradhan, S., Pellegrini, M., and Jacobsen, S. E. Conseration and divergence of methylation patterning in plants and animals. PNAS 2010;107:8689–8694.

5. Zemach, A., McDaniel, I. E., Silva, P., and Zilberman, D. Genome-wide evolutionary analysts of eukaryotic DNA methylation. Science 2010;328:916–919.

6. Urich, M. A., Nery, J. R., Lister, R., Schmitz, R. J., and Ecker, J. R. Methylcseq library preparation for base-resolution whole-genome bisulfite sequencing. Nature Protocols 2015;10:475–483.

7. Kuo, K. C., McCune, R. A., Gehrke, C. W., Midgett, R., Ehrlich, M. Quantitative reversed-phase high performance liquid chromatographic determination of major and modified deoxyribonucleosides in DNA. Nucleic Acids Research 1980;8:4763–4776.

8. Fraga, M. F., Uriol, E., Borja, D. L., Berdasco, M., Esteller, M., Cañal, M. J., Rodríguez, R. High-performance capillary electrophoretic method for the quantification of 5-methyl 2’-deoxycytidine in genomic DNA: application to plant, animal and human cancer tissues. Electrophoresis 2002;23:1677–1681.

9. Karimi M., Johansson S., Stach D., Corcoran M., Grander D. LUMA (LUminometric Methylation Assay)-a high throughput method to the analysis of genomic DNA methylation. Experimental Cell Research 2006;312:1989–1995.

10. Stroud, H., Greenberg, M. V. C., Feng, S., Bernatavichute, Y. V., and Jacobsen, S. E. Comprehensive analysis of silencing mutants reveals complex regulation comprehensive analysis of silencing mutants reveals complex regulation of the *Arabidopsis* methylome. Cell 2013; 152: 352–364.

11. Reik, W., Dean, W., and Walter, J. Epigenetic reprogramming in mammalian development. Science 2001;293:1089–1093.

12. Law, J. A., and Jacobsen, S. E. Establishing, maintaining and modifying DNA methylation patterns in plants and animals. Nature Reviews Genetics 2010;11:204–220

13. Popp, C., Dean, W., Feng, S., Cokus, S. J., Andrews, S., Pellegrini, M., Jacobsen, S. E., and Reik, W. Genome-wide erasure of DNA methylation in mouse primordial germ cells is affected by AID deficiency. Nature 2010;463:1101–1105.

14. Kobayashi, H., Sakurai, T., Imai, M., Takahashi, N., Fukuda, A., Yayoi, O., Sato, S., Nakabayashi, K., Hata, K., Sotomaru, Y., Suzuki, Y., and Kono, T. Contribution of intragenic DNA methylation in mouse gametic DNA methylomes to establish oocyte-specific heritable marks. PLoS Genetics 2012;8:e1002440

15. Seisenberger, S., Andrews, S., Krueger, F., Arand, J., Walter, J., Santos, F., Popp, C., Thienpont, B., Dean, W., Reik, W. The dynamics of genome-wide DNA methylation reprogramming in mouse primordial germ cells. Molecular Cell 2012;48:849–862.

16. Lister, R., Mukamel, E. A., Nery, J. R., Urich, M., Puddifoot, C. A., Johnson, N. D., Lucero, J., Huang, Y., Dwork, A. J., Schultz, M. D., Yu, M., Tonti-Filippini, J., Heyn, H., Hu, S., Wu, J. C., Rao, A., Esteller, M., He, C., Haghighi, F. G., Sejnowski, T. J., Behrens, M. M., and Ecker, J. R. Global epigenomic reconfiguration during mammalian brain development. Science 2013;341:1237905.

17. Hon, G. C., Rajagopal, N., Shen, Y., McCleary, D. F., Yue, F., Dang, M. Y., and Ren, B. Adult tissue methylomes harbor epigenetic memory at embryonic enhancers. Nature Genetics 2013;45:1198–1206.

18. Jackson, J. P., Lindroth, A. M., Cao X., and Jacobsen, S. E. Control of CpNpG DNA methylation by the KRYPTONITE histone H3 methyltransferase. Nature 2002;416:556–560.

19. Du, J., Zhong, X., Bernatavichute, Y.V., Stroud, H., Feng, S., Caro, E., Vashisht, A.A., Terragni, J., Chin, H.G., Tu, A., Hetzel, J., Wohlschlegel, J. A., Pradhan, S., Patel, D. J., and Jacobsen, S. E. Dual binding of chromomethylase domains to H3K9me2-containing nucleosomes directs DNA methylation in plants. Cell 2012;151:167–180.

20. Du, J., Johnson, L. M., Groth, M., Feng, S., Hale, C. J., Li, S., Vashisht, A. A., Gallego-Bartolome, J., Wohlschlegel, J. A., Patel, D. J., and Jacobsen, S. E. Mechanism of DNA methylation-directed histone methylation by KRYPTONITE. Molecular Cell 2014;55:495–504.

21. Calarco, J. P., Borges, F., Donoghue, M. T., Van Ex, F., Jullien, P. E., Lopes, T., Gardner, R., Berger, F., Feijo, J.A, Becker, J. D., and Martienssen, R. A. Reprogramming of DNA methylation in pollen guides epigenetic inheritance via small RNA. Cell 2012;151:194–205.

22. Schultz, M. D., He, Y., Whitaker, J. W., Hariharan, M., Mukamel, E. A., Leung, D., Rajagopal, N., Nery, J. R., Urich, M. A., Chen, H., Lin, S., Lin, Y., Jung, I., Schmitt, A. D., Selvaraj, S., Ren, B., Sejnowski, T. J., Wang. W., and Ecker, J. R. Human body epigenome maps reveal noncanonical DNA methylation variation. Nature 2015;523:212–216.

23. Martin, M. Cutadapt removes adapter sequences from high-throughput sequencing reads. EMBnet 2011;17:10–12.

24. Bolger, A. M., Lohse, M., and Usadel, B. Trimmomatic: a flexible trimmer for illumina sequence data. Bioinformatics 2014;30:2114–2120.

